# Two *Caenorhabditis elegans* calponin-related proteins have overlapping functions to maintain cytoskeletal integrity and are essential for reproduction

**DOI:** 10.1101/2020.04.29.069104

**Authors:** Shoichiro Ono, Kanako Ono

**Affiliations:** Department of Pathology, Department of Cell Biology, Winship Cancer Institute, Emory University School of Medicine, Atlanta, Georgia 30322

**Author notes:** To whom correspondence should be addressed: Department of Pathology, Emory University School of Medicine, 615 Michael Street, Whitehead Research Building, Room 105N, Atlanta, Georgia 30322.

**Keywords:** Actin, bundling, cytoskeleton, calponin-like (CLIK) motif, ovulation

## Abstract

Multicellular organisms have multiple genes encoding calponins and calponin-related proteins, and some of these are known to regulate actin cytoskeletal dynamics and contractility. However, functional similarities and differences among these proteins are largely unknown. In the nematode *Caenorhabditis elegans*, UNC-87 is a calponin-related protein with seven calponin-like (CLIK) motifs and is required for maintenance of contractile apparatuses in muscle cells. Here, we report that CLIK-1, another calponin-related protein that also contains seven CLIK motifs, has an overlapping function with UNC-87 to maintain actin cytoskeletal integrity *in vivo* and has both common and different actin-regulatory activities *in vitro*. CLIK-1 is predominantly expressed in the body wall muscle and somatic gonad, where UNC-87 is also expressed. *unc-87* mutation causes cytoskeletal defects in the body wall muscle and somatic gonad, whereas *clik-1* depletion alone causes no detectable phenotypes. However, simultaneous depletion of *clik-1* and *unc-87* caused sterility due to ovulation failure by severely affecting the contractile actin networks in the myoepithelial sheath of the somatic gonad. *In vitro*, UNC-87 bundles actin filaments. However, CLIK-1 binds to actin filaments without bundling them and is antagonistic to UNC-87 in filament bundling. UNC-87 and CLIK-1 share common functions to inhibit cofilin binding and allow tropomyosin binding to actin filaments, suggesting that both proteins stabilize actin filaments. Thus, partially redundant functions of UNC-87 and CLIK-1 in ovulation is likely mediated by their common actin-regulatory activities, but their distinct activities in actin bundling suggest that they also have different biological functions.

## Introduction

Calponins and calponin-related proteins regulate actin filament bundling, actin filament stability, and actomyosin contractility in a wide variety of cell types (1–3). Many of these proteins contain a single calponin-homology (CH) domain and various numbers of calponin-like (CLIK) motifs (also known as CLIK repeats). Although CH domains are recognized as actin-binding domains of several proteins, such as fimbrin, α-actinin, and spectrin (4,5), the CH domain of calponin is not an actin-binding site (6–9). Instead, CLIK motifs in the C-terminus of calponin act as actin-binding sites (6,9,10). CLIK motifs are 23-amino-acid sequences and present in variable numbers in calponins and calponin-related proteins (9,11). Vertebrate SM22/transgelin and yeast calponin-related proteins have only one CLIK motif (12–16). Vertebrate calponins have three CLIK motifs (17), whereas calponins in flatworms and mollusks have five CLIK motifs (18–20). UNC-87 in the nematode *Caenorhabditis elegans* has seven CLIK motifs (21). Importantly, the number of CLIK motifs correlates with the strength of actin-filament binding *in vitro* (22) and inhibitory effects on actin filament dynamics in cultured cells (10,23). These observations suggest that CLIK motifs are modular actin-binding sequences with incremental affinity with actin filaments depending on the number of motifs. However, whether all CLIK motifs have the same or different properties remains unknown.

Nematodes have uniquely evolved calponin-related proteins with multiple CLIK motifs and without a CH domain. *C. elegans* UNC-87, which contains seven CLIK motifs (21), is predominantly expressed in muscular tissues and associated with actin filaments (21,24,25). Two isoforms, UNC-87A and UNC-87B, are expressed by two alternative promoters for two separate first exons, and they share all CLIK motifs and similar biochemical properties (21,24). Mutations in the *unc-87* gene cause disorganization of sarcomeric actin filaments in the body wall muscle due to defects in the maintenance of sarcomeres (21,26,27). UNC-87 is conserved in other nematode species and play a critical role in worm motility (28,29). *In vitro*, UNC-87 bundles actin filaments (22), inhibits actin-filament severing by actin-depolymerizing factor (ADF)/cofilin (25), and inhibits actomyosin ATPase (24). In addition, the *C. elegans* genome has three other genes encoding similar calponin-related proteins with various numbers of CLIK motifs: *clik-1* (seven motifs), *clik-2* (six motifs), and *clik-3* (five motifs). A proteomic study reported that CLIK-1, CLIK-2, and CLIK-3 are enriched in the pharyngeal muscle (30). Null mutations of *clik-2* and *clik-3* cause pumping defects in the pharynx, whereas null mutation of *clik-1* causes no detectable phenotypes (31). Therefore, function of *clik-1* remains unknown. In this study, we aimed to determine the function of *clik-1*. Our results indicate that UNC-87 and CLIK-1 are coexpressed in muscle tissues and have overlapping and critical functions in the reproductive system by maintaining cytoskeletal integrity in the somatic gonad. Biochemical characterization showed that UNC-87 and CLIK-1 have common and distinct actin-regulatory activities *in vitro*.

## Results

### CLIK-1 and UNC-87 are partially redundant and required for C. elegans reproduction

*C. elegans* has four genes (*unc-87, clik-1, clik-2*, and *clik-3*) encoding calponin-related proteins with no CH domains. Among them, primary structures of UNC-87B (hereafter referred to as UNC-87) and CLIK-1 (also known as T25F10.6, GenBank accession number: CCD74240) are very similar (42 % identity) containing seven CLIK motifs (C1 - C7) (Fig. 1A). Comparison of CLIK motifs from representative calponins and calponin-related proteins shows that CLIK motifs are highly conserved across species (Fig. 1B). However, sequences of C5, C6, and C7 of CLIK-1 are diverged in their N-terminal halves (Fig. 1B), suggesting that biochemical properties of CLIK-1 are somewhat different from those of UNC-87.

**Figure 1.**
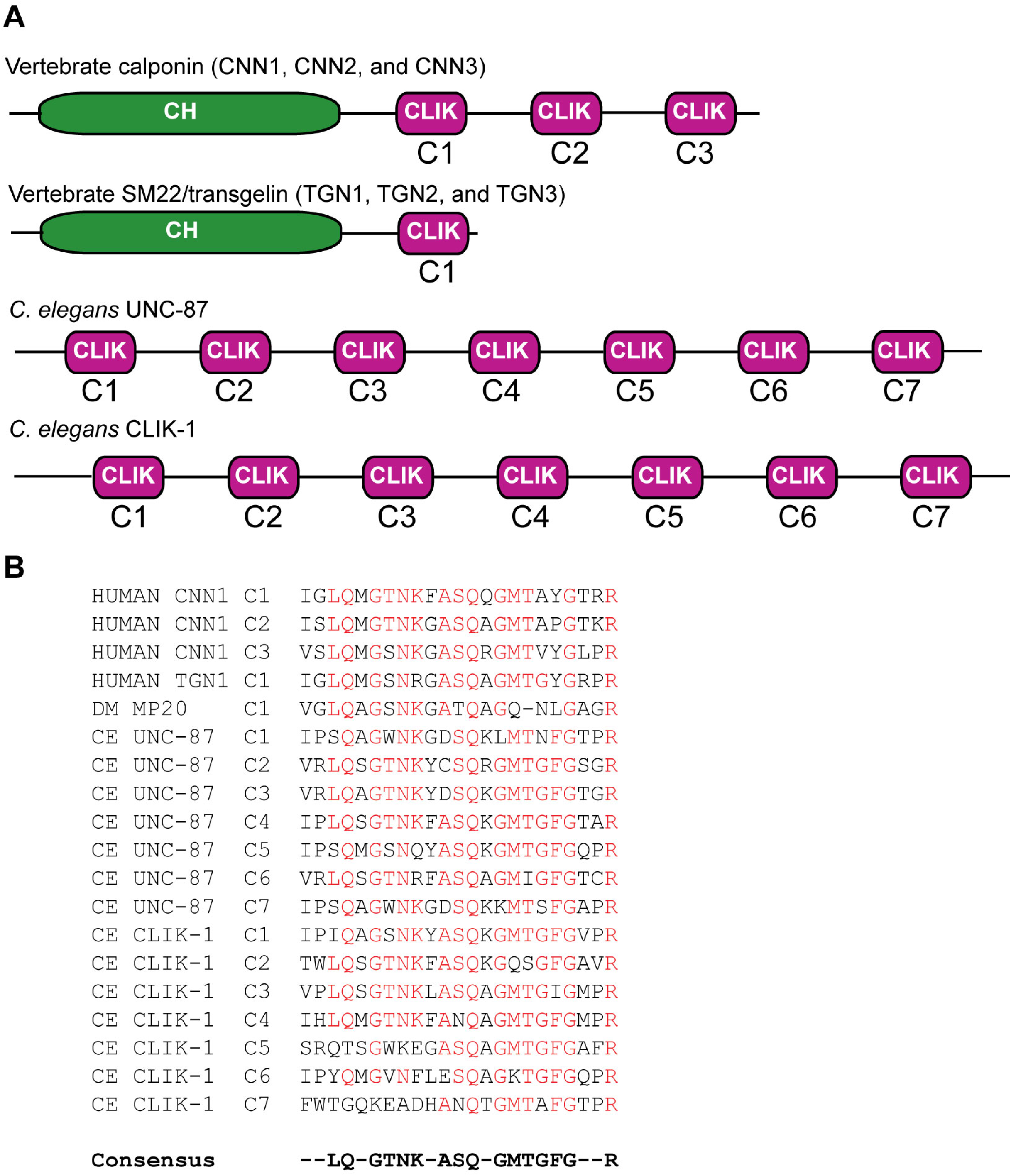
Domain organizations of calponin and calponin-related proteins. (A) Schematic representation of domain organizations of vertebrate calponin (CNN1, calponin 1; CNN2, calponin 2; CNN3, calponin 3), vertebrate SM22/transgelin (TGN1, SM22α/transgelin 1; TGN2, transgelin 2; TGN3, transgelin 3), *C. elegans* UNC-87, and *C. elegans* CLIK-1. CH: calponin-homology domain; CLIK: calponin-like motif. (B) Sequence alignment of CLIK motifs from calponin and calponin-related proteins. Human calponin 1 (CNN1) (NCBI Reference Sequence: NM_001299); human SM22α/transgelin (TGN1) (NCBI Reference Sequence: NM_001001522); *Drosophila melanogaster* MP20 (NCBI Reference Sequence: NM_057295); *C. elegans* UNC-87 (NCBI Reference Sequence: NM_001025922); *C. elegans* CLIK-1 (GenBank accession number: CCD74240).

To determine expression pattern and subcellular localization of CLIK-1, green fluorescent protein (GFP) was fused to the C-terminus of CLIK-1 at the endogenous *clik-1* locus using CRISPR/Cas9-mediated genome editing. CLIK-1-GFP was strongly expressed in the body wall muscle (Fig. 2A, C, E) and myoepithelial sheath of the somatic gonad (Fig. 2B, D, F), where CLIK-1-GFP localized in a sarcomeric pattern in the body wall muscle (Fig. 2A) and in a filamentous pattern in the myoepithelial sheath (Fig. 2B). Our preliminary study indicated that CLIK-1 co-localized with actin filaments (our unpublished observastions). However, we did not detect expression of CLIK-1-GFP in the pharynx, although we did detect expression of CLIK-1-GFP in the neurons surrounding the pharynx (our unpublished observations). Thus, our results disagree with the proteomics study reporting enrichment of CLIK-1 in the pharynx (30). Body wall muscle and myoepithelial sheath of the gonad are tissues where UNC-87 is also expressed and associated with actin filaments (21,24,25), suggesting that CLIK-1 and UNC-87 regulate actin filaments in the same cells. Additional characterization of tissue distribution and subcellular localization of CLIK-1-GFP will be reported elsewhere.

**Figure 2.**
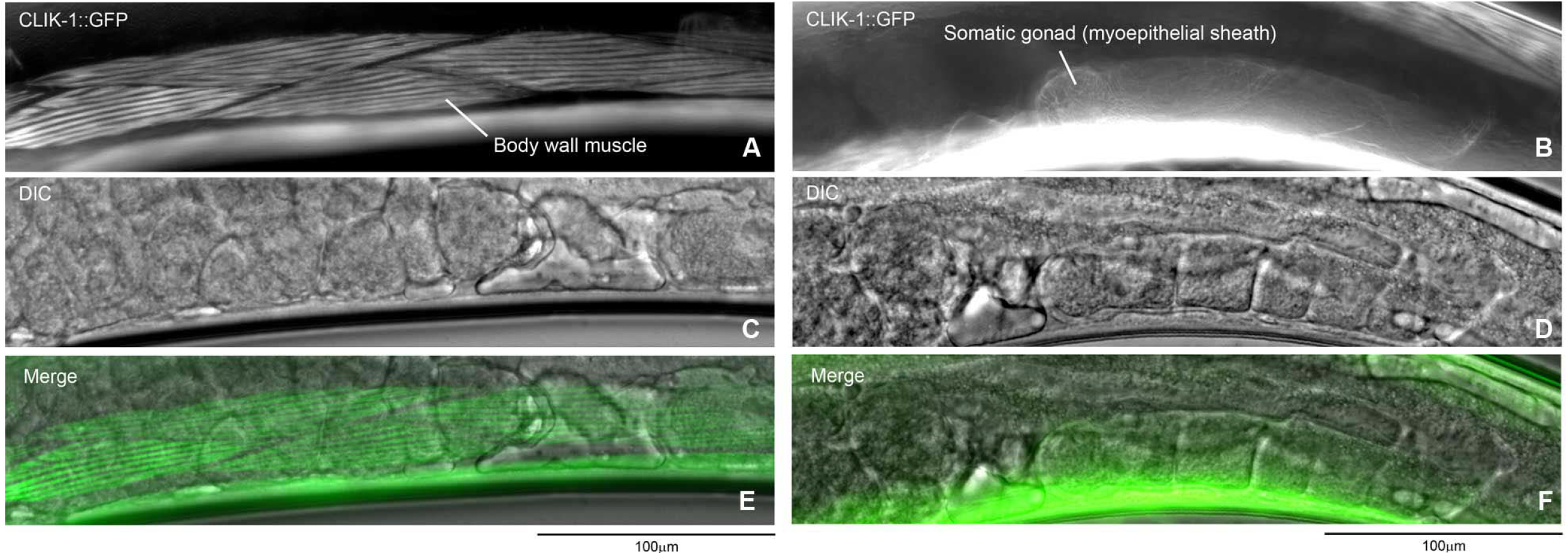
Expression and localization of CLIK-1 in *C. elegans*. GFP was fused to the C-terminus of CLIK-1 using CRISPR/Cas9-mediated genome editing. Localization of CLIK-1-GFP (A, B) and differential interference contrast images (C, D) were examined in live adult hermaphrodites. Merged images are shown in E and F. Expression of CLIK-1-GFP in the body wall muscle (A, C, E) and the myoepithelial sheath of the somatic gonad (B, D, F) are highlighted.

In a recent study, Wang et al. reported that *clik-1* gene knockout did not cause any obvious phenotypes (31), and we confirmed their observations using a separately isolated *clik-1*-null strain [*clik-1(ok2355)*] that contains a 1-kb deletion in the *clik-1* gene (our unpublished observation). However, we found that knockdown of *clik-1* by RNA interference (RNAi) in an *unc-87* null mutant [*unc-87(e1459)*] (21,24) caused sterility with severe cytoskeletal defects in the reproductive system (Fig. 3 and 4). When worms were treated with control RNAi, wild-type worms actively moved and produced many progeny (Fig. 3A and I), whereas *unc-87(e1459)* worms were slow-moving and produced much less progeny (Fig. 3C and I). RNA interference of *clik-1* did not affect movement and fecundity in wild-type background (Fig. 3B and I), but enhanced impaired worm motility to nearly paralysis (Fig. 3D and I) and caused sterility in *unc-87(e1459)* (Fig. 3D, see the absence of eggs and small worms on the culture plate). In the body wall muscle, actin was organized into a striated sarcomeric pattern in wild-type with control RNAi (Fig. 3E) or *clik-1(RNAi)* (Fig. 3F). Sarcomeric actin in *unc-87(e1459)* with control RNAi was disorganized with accumulation in aggregates (Fig. 3G), and the actin disorganization was not apparently enhanced by *clik-1(RNAi)* in *unc-87(e1459)* (Fig. 3H), suggesting that enhancement of paralysis by *clik-1 (RNAi)* is not due to structural defects in the sarcomeres. Currently, we do not know why motility of *unc-87(e1459); clik-1(RNAi)* worms are severely impaired.

**Figure 3.**
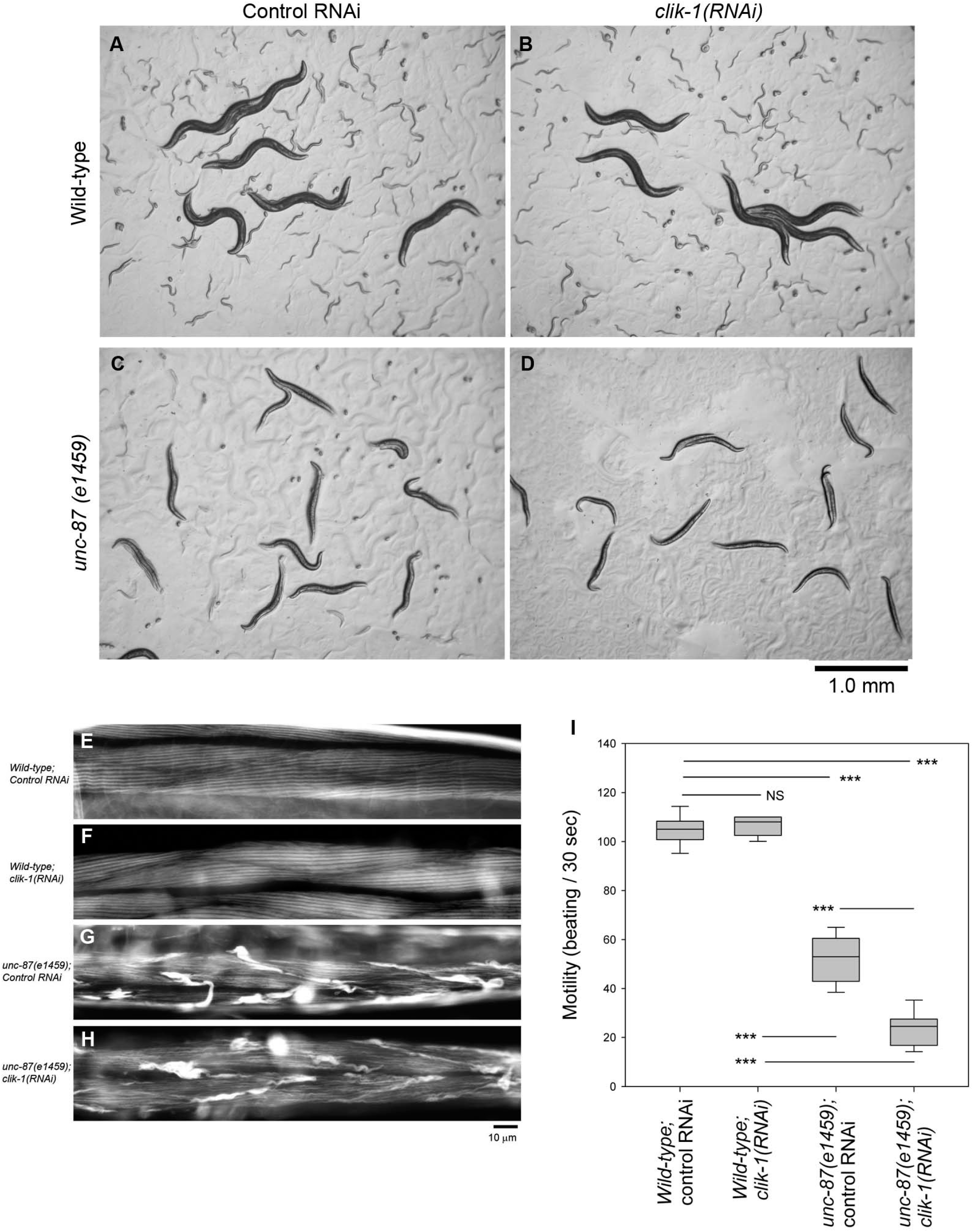
Simultaneous depletion of *unc-87* and *clik-1* causes impaired worm motility and sterility. Wild-type or *unc-87(e1459)* worms were treated with control RNAi or *clik-1(RNAi)*. (A – D) Live worms on the culture plates are shown. Large worms are the RNAi-treated worms. Small worms and eggs are F1 progeny from the treated worms. Bar, 1.0 mm. (E-H) Organization of actin filaments in the body wall muscle as observed by staining with tetramethylrhodamine-phalloidin. Bar, 10 μm. (I) Worm motility was quantified as number of beating per 30 sec. n = 10. Boxes represent the range of the 25th and 75th percentiles, with the medians marked by solid horizontal lines, and whiskers indicate the 10th and 90th percentiles. NS, not significant. ***, p < 0.001.

**Figure 4.**
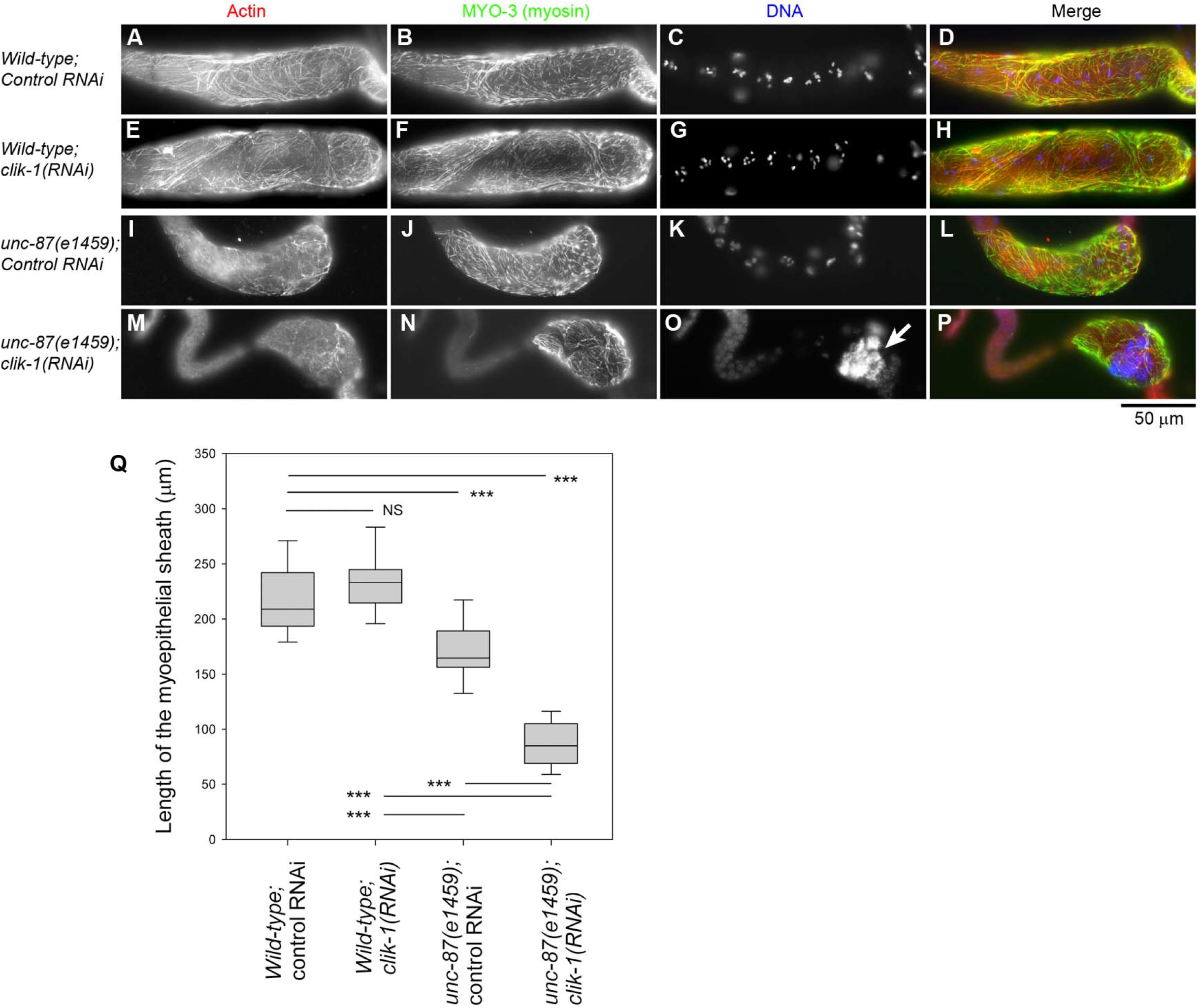
Simultaneous depletion of *unc-87* and *clik-1* causes severe disorganization of actin filaments in the somatic gonad. Wild-type or *unc-87(e1459)* worms were treated with control RNAi or *clik-1(RNAi)*. (A-P) Dissected gonads were stained for F-actin (A, E, I, M), MYO-3 (myosin) (B, F, J, N), and DNA (C, G, K, O). The proximal side of the gonads are oriented to the right. Merged imaged (F-actin in red, MYO-3 in green, and DNA in blue) are shown in the right column (D, H, L, P). Bar, 50 μm. (Q) Lengths of the myoepithelial sheath as marked by the MYO-3 staining were measured. Sample sizes (n) were 29 (wild-type; control RNAi), 28 (wild-type; *clik-1(RNAi)*), 26 (*unc-87(e1459);* control RNAi), and 27 (*unc-87(e1459); clik-1(RNAi)*). Boxes represent the range of the 25th and 75th percentiles, with the medians marked by solid horizontal lines, and whiskers indicate the 10th and 90th percentiles. NS, not significant. ***, p < 0.001.

Simultaneous depletion of *unc-87* and *clik-1* disrupted cytoskeletal integrity in the somatic gonads, which explains why worm reproduction was strongly impaired (Fig. 4). In *C. elegans* hermaphrodites, oocytes are surrounded by the myoepithelial sheath in the gonad that provides contractile forces during ovulation of mature oocytes (32,33). The myoepithelial sheath is a smooth-muscle-like somatic tissue that contains non-striated actomyosin networks (34–36). The MYO-3 myosin heavy chain was used as a marker for the myoepithelial sheath (34,36,37). In wild-type with control RNAi, the myoepithelial sheath contained networks of actin and MYO-3 myosin and covered multiple oocytes (as shown by clusters of condensed chromosomes) (Fig. 4A-D). RNAi of *clik-1* in wild-type did not cause alteration in the actomyosin networks in the myoepithelial sheath (Fig. 4E-H). In *unc-87(e1459)* with control RNAi, actin filaments were somewhat disorganized, and the myoepithelial sheath was not as extended as wild-type with control RNAi (Fig. 4I-L and Q). Strikingly, in *unc-87(e1459)* with *clik-1(RNAi)*, actin filaments were only sparsely distributed, and the myoepithelial sheath was shortened and nearly collapsed to one side of the gonad (Fig. 4M-P and Q). The oocytes in these worms contained large accumulations of DNA (Fig. 4O, arrow) as a result of endomitotic DNA replication during ovulation failure (38). These phenotypes indicate that UNC-87 and CLIK-1 have partially redundant function to maintain actin cytoskeletal integrity in the myoepithelial sheath and to execute proper ovulation that is an essential process in worm reproduction.

### CLIK-1 competes with UNC-87 for actin filament binding and inhibits filament bundling

To characterize biochemical properties of CLIK-1 and compare them with those of UNC-87, we prepared recombinant CLIK-1 protein with no extra tag sequence (Fig. 5A). When CLIK-1 was incubated with filamentous (F-) actin, CLIK-1 co-sedimented with F-actin after ultracentrifugation (Fig. 5B) indicating that CLIK-1 bound to F-actin. Quantitative analysis of the F-actin co-sedimentation assays at various CLIK-1 concentrations demonstrated that CLIK-1 bound to F-actin with a dissociation constant (*Kd*) of 0.51 ± 0.094 μM (n = 3), which is higher than the estimated *Kd* (~0.15 μM) for UNC-87 binding to F-actin (25). Binding was saturated at ~0.25 (mol CLIK-1/mol actin) indicating that CLIK-1 binds to actin filaments with a stoichiometry of 1:4 (Fig. 5C). Under low-speed centrifugation (18,000 x g, 10 min), CLIK-1 did not increase actin in the pellets (Fig. 5B and D), whereas UNC-87 increased actin in the pellets by bundling filaments (22) (Fig. 5D). Therefore, CLIK-1 binds to actin filaments but, unlike UNC-87, lacks filament bundling activity.

**Figure 5.**
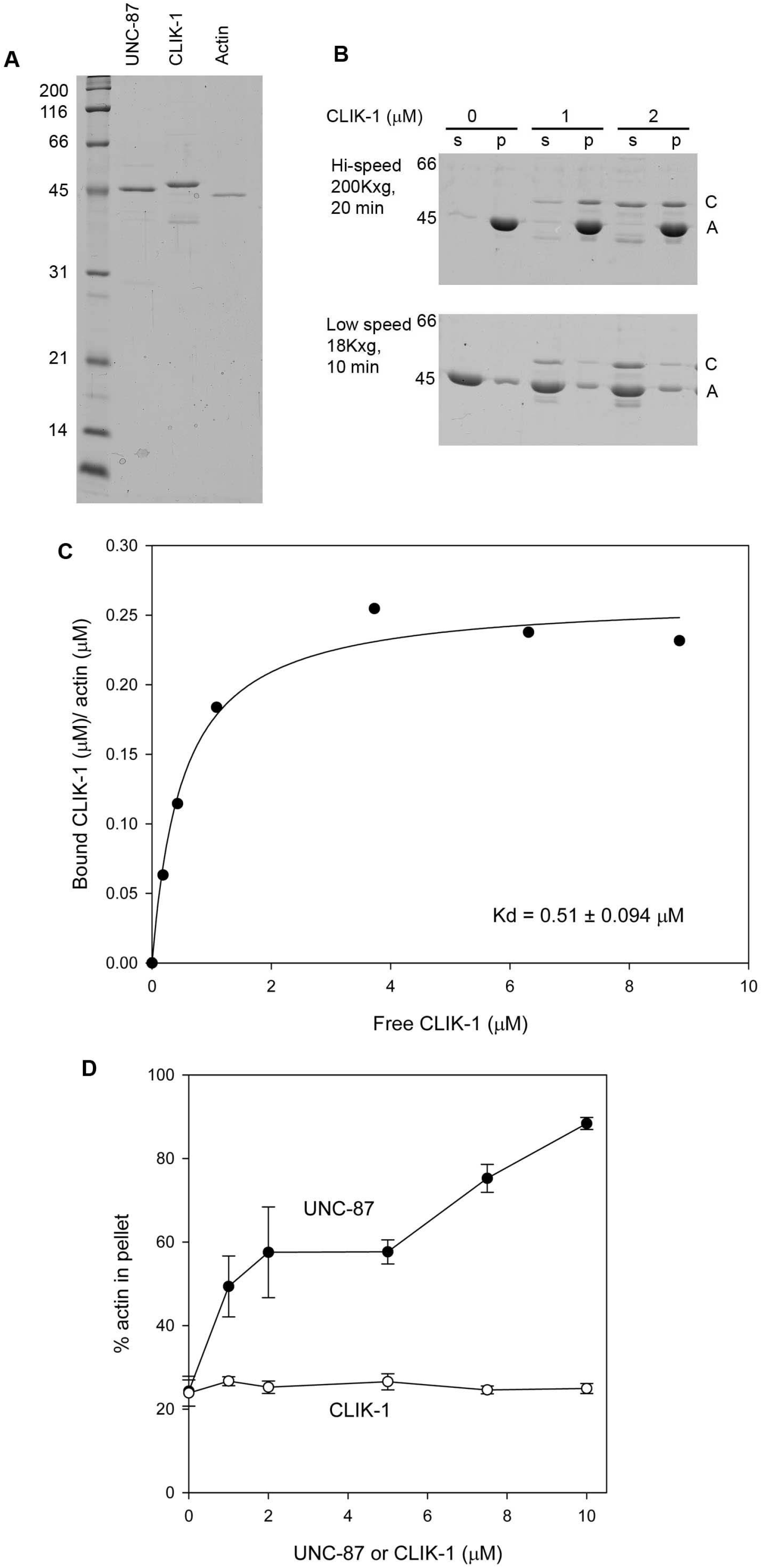
CLIK-1 binds to actin filaments without bundling them. (A) Purity of recombinant UNC-87 (0.5 μg), recombinant CLIK-1 (0.5 μg), and rabbit muscle actin (0.2 μg), was examined by SDS-PAGE (12 % acrylamide gel). Molecular weight markers in kDa are shown on the left. (B) F-actin sedimentation assays at high speed (200,000 x g, 20 min) (top) and low speed (18,000 x g, 10 min) (bottom). F-actin (10 μM) was incubated with 0 – 2 μM CLIK-1 for 1 hr and centrifuged at the indicated conditions. Supernatants (s) and pellets (p) were separated and examined by SDS-PAGE. Positions of CLIK-1 (C) and actin (A) are indicated on the right. (C) Quantitative analysis of the high-speed F-actin sedimentation assays. The high-speed F-actin sedimentation assays were performed using 5 μM F-actin and 0 – 10 μM CLIK-1. Molar ratios of CLIK-1 to actin in the pellets (actin-dependent sedimentation of CLIK-1 was determined by subtracting non-specific sedimentation of CLIK-1) [bound CLIK-1 (μM)/actin (μM)] are plotted as a function of free CLIK-1. A dissociation constant (*Kd*) was determined from three independent experiments (means ± standard deviation). (D) Quantitative analysis of the low-speed F-actin sedimentation assays. The low-speed F-actin sedimentation assays were performed with 10 μM F-actin and 0 – 10 μM UNC-87 or CLIK-1. Percentages of actin in the pellets (bundled actin filaments) were plotted as a function of concentration of UNC-87 or CLIK-1. Data are means ± standard deviation from three independent experiments.

Because CLIK-1 and UNC-87 localize to actin filaments in the same cells, we examined their functional relationship when the two proteins are both present. Not surprisingly from their sequence similarity, CLIK-1 and UNC-87 competed for F-actin binding (Fig. 6A-F). In the F-actin co-sedimentation assays at high speed (200,000 x g, 20 min) in the presence of a constant UNC-87 concentration, increasing concentrations of CLIK-1 reduced UNC-87 in the pellets (Fig. 6A and B). In the opposite experiments with a constant CLIK-1 concentration, increasing concentrations of UNC-87 reduced CLIK-1 in the pellets (Fig. 6C and D). UNC-87 was more resistant to dissociation than CLIK-1 and more effective to dissociate the counterpart than CLIK-1, which is consistent with the higher affinity of UNC-87 with actin than that of CLIK-1. Interestingly, the competition between UNC-87 and CLIK-1 resulted in antagonistic modulation of actin filament bundling. UNC-87 bundled actin filaments and increased actin in the pellets under low-speed centrifugation (Fig. 6E and F). However, increasing concentrations of CLIK-1 decreased actin in the pellets even in the presence of UNC-87 (Fig. 6E and F). Their antagonistic effects on actin bundling were further demonstrated by direct observation of fluorescently labeled actin filaments. Actin filament bundles were formed in the presence of UNC-87 (Fig. 6, compare G and H), but not CLIK-1 (Fig. 6I). When both UNC-87 (1 μM) and CLIK-1 (2 μM) were present, very few actin bundles were formed (Fig. 6J and K). The concentration of CLIK-1 needed to be higher than that of UNC-87 to cause significant reduction of actin bundles (Fig. 6K). Thus, the balance of UNC-87 and CLIK-1 can determine the extent of actin filament bundling *in vitro*.

**Figure 6.**
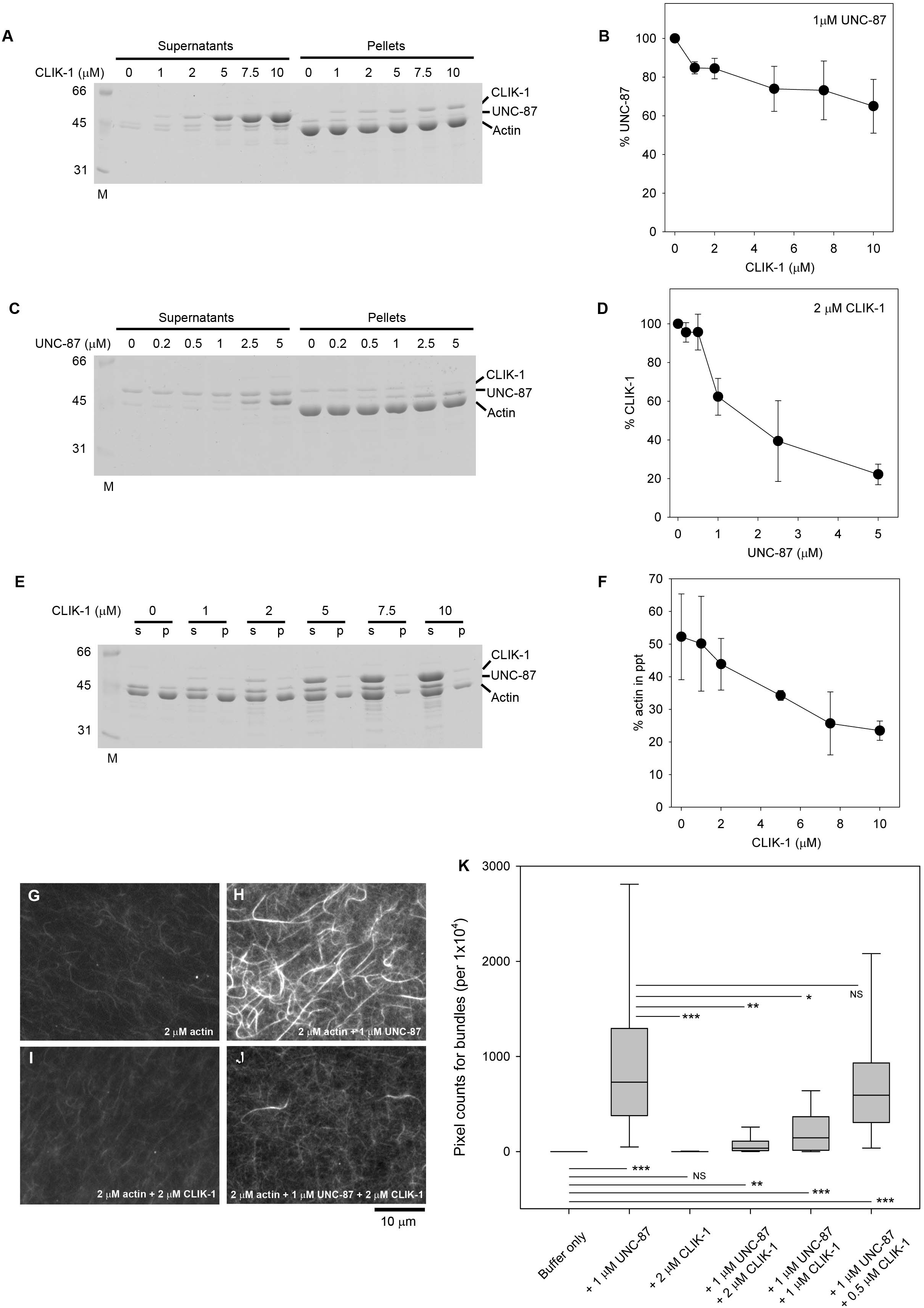
Competitive binding of UNC-87 and CLIK-1 to actin filaments. (A, B) F-actin (10 μM) was incubated with 1 μM UNC-87 and 0 – 10 μM CLIK-1 and examined by high-speed co-sedimentation assays. Supernatants and pellets were analyzed by SDS-PAGE (A). Relative amounts of UNC-87 in the pellets (100 % in the absence of CLIK-1) were quantified and plotted as a function of CLIK-1 concentration (B). Data are means ± standard deviation from three independent experiments. (C, D) Factin (10 μM) was incubated with 2 μM CLIK-1 and 0 – 5 μM UNC-87 and examined by high-speed cosedimentation assays. Supernatants and pellets were analyzed by SDS-PAGE (C). Relative amounts of CLIK-1 in the pellets (100 % in the absence of UNC-87) were quantified and plotted as a function of UNC-87 concentration (D). Data are means ± standard deviation from three independent experiments. (E, F) F-actin (10 μM) was incubated with 2.5 μM UNC-87 and 0 – 10 μM CLIK-1 and examined by lowspeed sedimentation assays. Supernatants (s) and pellets (p) were analyzed by SDS-PAGE (E). Percentages of actin in the pellets (bundled actin filaments) were plotted as a function of CLIK-1 concentration. Data are means ± standard deviation from four independent experiments. (G – K) Direct observation of actin bundling by fluorescence microscopy. DyLight 549-labeled actin (2 μM) was incubated with buffer only (G) or buffer with 1 μM UNC-87 (H), 2 μM CLIK-1 (I), or 1 μM UNC-87 and 2 μM CLIK-1 (J) and observed with fluorescence microscopy. Bar, 10 μm. Actin bundle formation was quantified (K) from randomly selected regions of interest (100 x 100 pixels) (n = 25). Boxes represent the range of the 25th and 75th percentiles, with the medians marked by solid horizontal lines, and whiskers indicate the 10th and 90th percentiles. NS, not significant. *, 0.01 < p < 0.05; **, 0.001 < p < 0.01; ***, p < 0.001.

### Tropomyosin binds to actin filaments simultaneously with CLIK-1 or UNC-87 and partially inhibits filament bundling

Although the biochemical experiments suggest that CLIK-1 is an inhibitor of actin filament bundling, we did not detect excessive actin filament bundling in the body wall muscle or myoepithelial sheath of *clik-1(RNAi)* worms or *clik-1* knockout worms (our unpublished observation). Therefore, we reasoned that other actin binding protein(s) can inhibit actin filament bundling by UNC-87. Since tropomyosin is a major F-actin binding protein, we examined how tropomyosin interacts with actin filaments in the presence of UNC-87 or CLIK-1. The *lev-11* gene encodes at least six tropomyosin isoforms, and LEV-11A is a major high-molecular-weight isoform in muscle cells (39–42). LEV-11A bound to actin filaments in the presence of either UNC-87 or CLIK-1 (Fig. 7A-C). In addition, in the presence of a saturating concentration of LEV-11A, actin bundling, as determined by low-speed actin sedimentation, was inhibited when the UNC-87 concentrations were low (Fig. 7D-F). At high UNC-87 concentrations, actin filament bundling was no longer inhibited by LEV-11A (Fig. 7D-F). Therefore, tropomyosin, in addition to CLIK-1, can also inhibit UNC-87-induced actin filament bundling under certain conditions.

**Figure 7.**
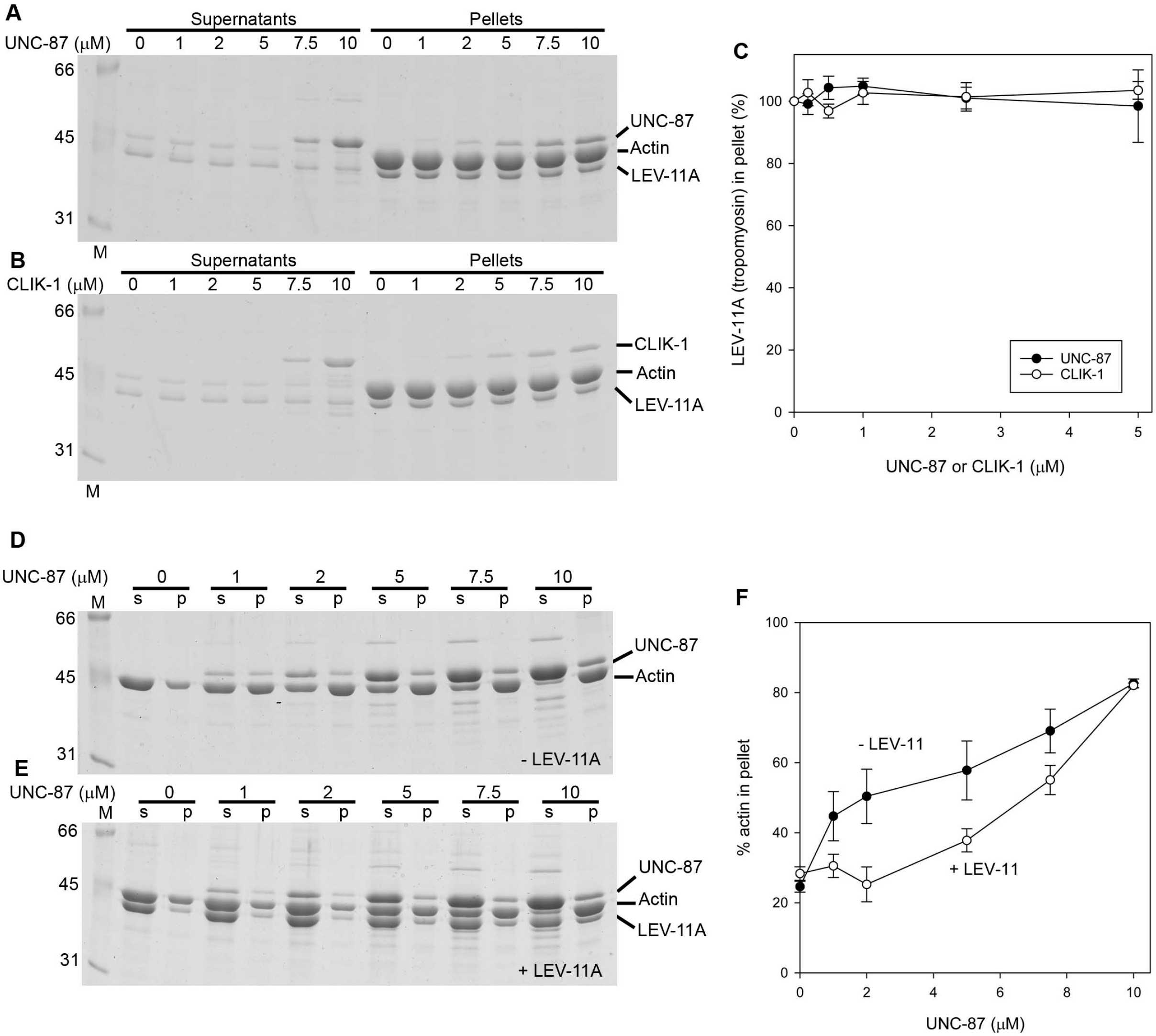
Tropomyosin binds to F-actin in the presence of UNC-87 or CLIK-1 and partially inhibits UNC-87-mediated actin filament bundling. (A – C) F-actin (10 μM) was incubated with 1 μM LEV-11A (tropomyosin) and 0 – 10 μM UNC-87 (A) or CLIK-1 (B) and examined by high-speed cosedimentation assays. Supernatants and pellets were analyzed by SDS-PAGE. Relative amounts of LEV-11A in the pellets (100 % in the absence of UNC-87 or CLIK-1) were quantified and plotted as a function of UNC-87 or CLIK-1 concentration (C). Data are means ± standard deviation from three independent experiments. (D – F) F-actin (10 μM) was incubated with 0 – 10 μM UNC-87 in the absence (D) or presence of 2 μM LEV-11A (E) and examined by low-speed sedimentation assays. Supernatants (s) and pellets (p) were analyzed by SDS-PAGE. Percentages of actin in the pellets (bundled actin filaments) were plotted as a function of UNC-87 concentration. Data are means ± standard deviation from three independent experiments.

### CLIK-1 and UNC-87 prevent ADF/cofilin from F-actin binding

UNC-87 stabilizes actin filaments by preventing biding of actin depolymerizing factor (ADF)/cofilin to the filaments (25). Similarly, CLIK-1 inhibited ADF/cofilin binding to actin filaments. UNC-60B, a muscle-specific ADF/cofilin isoform in *C. elegans*, severs actin filaments, and UNC-60B-bound actin filaments can be sedimented by ultracentrifugation (43–46). UNC-60B, which co-sedimented with Factin, decreased when increasing concentrations of UNC-87 (Fig. 8A and C) or CLIK-1 (Fig. 8B and C) were included. UNC-87 was more efficient to prevent actin binding of UNC-60B than CLIK-1 (Fig. 8C), which is consistent with the difference in their affinity with actin. Thus, UNC-87 and CLIK-1 share a common actin-stabilizing function by antagonizing ADF/cofilin.

**Figure 8.**
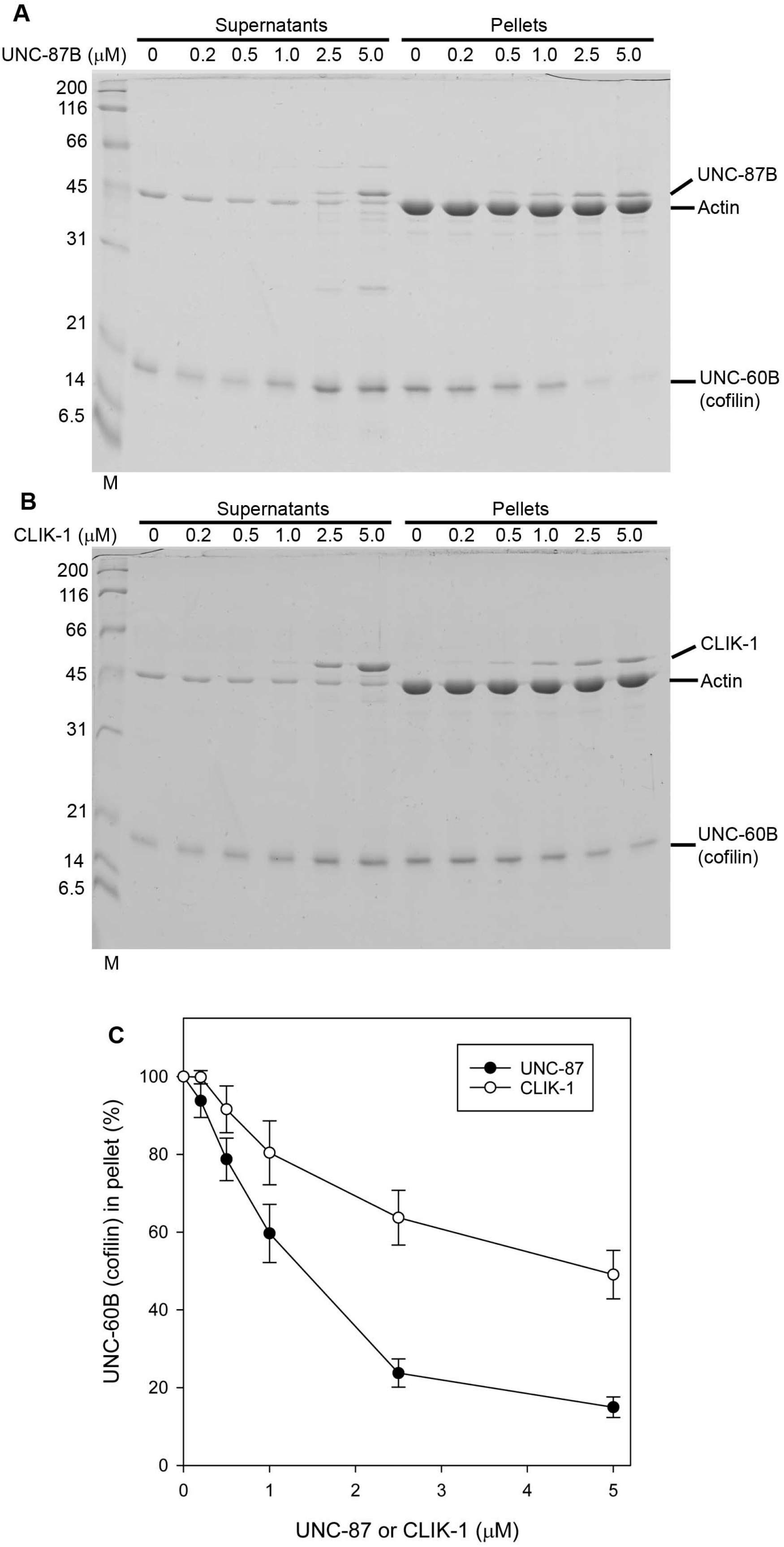
UNC-87 and CLIK-1 inhibit F-actin binding of ADF/cofilin. F-actin (10 μM) was incubated with 10 μM UNC-60B (ADF/cofilin) and 0 - 5 μM UNC-87 (A) or CLIK-1 (B) and examined by highspeed co-sedimentation assays. Supernatants and pellets were analyzed by SDS-PAGE. Relative amounts of UNC-60B in the pellets (100 % in the absence of UNC-87 or CLIK-1) were quantified and plotted as a function of UNC-87 or CLIK-1 concentration (C). Data are means ± standard deviation from three independent experiments.

## Discussion

Here, we demonstrate that two calponin-related actin binding proteins, UNC-87 and CLIK-1, are critical for cytoskeletal integrity in the *C. elegans* reproductive system. Depletion of both UNC-87 and CLIK-1 cause severe disorganization of actin filaments in the myoepithelial sheath of the somatic gonad, which results in sterility. Single depletion of either UNC-87 or CLIK-1 caused much milder or no phenotypes, indicating that UNC-87 and CLIK-1 have partially redundant functions *in vivo*. A previous single-cell proteomic study reported that CLIK-1 is enriched in the pharyngeal muscle cells (30). However, we do not detect strong expression of CLIK-1 in the pharynx. Instead, CLIK-1 expression is strong in other contractile tissues, including body wall muscle and myoepithelial sheath and overlaps with UNC-87 expression. Interestingly, biochemical analysis reveals both common and distinct actin-regulatory properties of UNC-87 and CLIK-1. Both proteins bind to actin filaments and prevent binding of ADF/cofilin to actin filaments. However, UNC-87 bundles actin filaments, but CLIK-1 does not. Their competitive binding to actin filaments modulates the extent of filament bundling when the two proteins co-exist. Thus, our current study uncovers novel functional aspects of calponin-related proteins in the regulation of the actin cytoskeleton.

The myoepithelial sheath of the somatic gonad (ovary) is highly sensitive to depletion of UNC-87 and CLIK-1. The myoepithelial sheath is a monolayer of smooth-muscle-like cells that contain non-striated contractile actin networks (34–36). Its contraction is tightly coupled with oocyte maturation and pushes only one mature oocyte out of the ovary for subsequent fertilization (32,33). Contraction of the myoepithelial sheath is regulated by both troponin-tropomyosin complex (47–49) and phosphorylation of myosin light chain (37). UNC-87 has an inhibitory effect on actomyosin contractility, and may facilitate relaxation of the myoepithelial sheath (24). Severe disorganization of the actin filaments in the myoepithelial sheath in double depletion of UNC-87 and CLIK-1 suggest that the two calponin-related proteins have redundant roles to stabilize actin filaments. CLIK-1 shares a common function with UNC-87 to protect actin filaments from ADF/cofilin (Fig. 8). ADF/cofilin and actin-interacting protein 1 are required for assembly of actin networks in the myoepithelial sheath (50,51). However, excessive activities of these proteins are expected to destabilize actin filaments. Either one of UNC-87 or CLIK-1 is sufficient to stabilize actin filaments, but loss of both proteins can cause severe actin disorganization such that the myoepithelial sheath is no longer functional. In addition, ovulatory contraction of the myoepithelial sheath involves extensive morphological changes and mechanical stress (32,33), and these calponin-related proteins may have a role in providing mechanical stability to the actin filaments as demonstrated biophysically for mammalian calponin (52,53).

To our knowledge, CLIK-1 is the only calponin-related protein lacking actin filament bundling activity. Because only a limited number of calponin-related proteins have been characterized biochemically, there may be other calponin-related proteins with variable functions. Vertebrate calponin bundles actin filaments *in vitro* (54,55) and play important roles in cytoskeletal structures containing actin bundles (10,56). Vertebrate SM22/transgelin (57,58) and yeast calponin-related proteins (12–14,59) have only one CLIK motif and still bundles actin filaments *in vitro*. A calponin-related protein with no actin bundling activity, such as CLIK-1 or an equivalent protein, may compete with conventional calponin and/or SM22/transgelin to modulate the extent of actin bundle formation. Although such an antagonistic relationship was not clearly detected between UNC-87 and CLIK-1 in the body wall muscle and myoepithelial sheath *in vivo*, it may be of functional significance in other cell types, or other actin-binding proteins, such as tropomyosin, may participate in the modulation of actin bundle formation.

The differences in the actin-regulatory activities of UNC-87 and CLIK-1 suggest that not all CLIK motifs behave in the same manner. Previous studies suggested that CLIK motifs are functionally similar modular motifs because the number of CLIK motifs correlates with the effectiveness in stabilizing actin filaments in cultured cells (10,23). The difference between UNC-87 and CLIK-1 in actin bundling activity indicate that some of the CLIK motifs in UNC-87 specifically mediate actin bundling. Since UNC-87 is monomeric (22), a single UNC-87 molecule needs to bind to multiple actin filaments to bundle them, suggesting that some of the CLIK motifs preferentially bind to a second filament. By contrast, CLIK-1 does not bundle actin filaments. Therefore, all the CLIK motifs in CLIK-1 bind to the same filament. There may be a difference in cooperativity among actin binding sites. In UNC-87, a first actin-binding site may prevent a second actin-binding site from binding to the same filament, while, in CLIK-1, a first actin-binding site may promote binding of a second actin-binding site to the same filament. Alternatively, the structure of UNC-87 may be suited to cross-bridge multiple filaments, whereas that of CLIK-1 may be optimal to bind to a single filament. Further dissection of biochemical properties of CLIK motifs should inform us about how different CLIK motifs contribute to functional differentiation of calponin-related proteins.

## Experimental Procedures

### C. elegans strains and culture

Worms were cultured following standard methods (60). The following strains were obtained from the *Caenorhabditis* Genetics Center and used in this study: N2 wild type, CB1459 *unc-87(e1459)*, and RB1820 *clik-1(ok2355). clik-1(ok2355)* was outcrossed three times and used in this study (outcrossed strain: ON225).

### CRISPR/Cas9 genome editing

The GFP sequence was inserted in the genome in-frame at the 3’-end of the *clik-1* coding region by CRISPR/Cas9-mediated genome editing using vectors and protocols described by Dickinson et al. (61). A single guide (sg) RNA target sequence (GATGTAATTACCACATACGA) was cloned into pDD162 (Addgene Plasmid # 47549) for expression of both sgRNA and Cas9 nuclease. Homology arms of ~500 bp each were fused with the GFP::self-excising cassette region from pDD282 (Addgene Plasmid #66823) by fusion PCR using Q5 High-Fidelity DNA Polymerase (New England Biolabs) and used as a homologous repair template. Hygromycin-resistant worms with a roller phenotype were isolated as knock-in worms. They were treated at 34 °C for 4 hours to induce excision of the self-excising cassette. In the next generation, non-roller GFP-positive worms were isolated, and three independent *clik-1::GFP* strains, ON352 *clik-1(kt1 [clik-1::gfp])*, ON353 *clik-1(kt2 [clik-1::gfp])*, and ON354 *clik-1(kt3 [clik-1::gfp])*, were established.

### RNA interference experiments

RNA interference experiments were performed by feeding with *Escherichia coli* HT115(DE3) expressing double-stranded RNA as described previously (62). The RNAi experiments were started by treating L1 larvae, and phenotypes were observed when they grew to adult worms after three days. Control RNAi experiments were performed using the unmodified L4440 plasmid vector (kindly provided by Andrew Fire, Stanford University) (63). An RNAi clone for *clik-1* (V-4N16) was obtained from Source BioScience. Images of worms on agar plates were captured using a Nikon Coolpix 995 digital camera mounted on a Zeiss Stemi 2000 stereo microscope.

### Fluorescence microscopy

Live worms were anesthetized in M9 buffer containing 0.1% tricaine and 0.01% tetramisole for 30 min, mounted on 2% agarose pads, and observed by epifluorescence and differential interference contrast optics using a Nikon Eclipse TE2000 inverted microscope (Nikon Instruments, Tokyo, Japan).

Staining of whole worms with tetramethylrhodamine–phalloidin (Sigma-Aldrich) was performed as described previously (64).

Gonads were dissected from adult hermaphrodites on polylysine-coated glass slides as described previously (36). They were fixed by methanol at −20°C for 5 min and washed by phosphate-buffered saline (PBS) and primary antibodies in PBS containing 1% BSA. After being washed with PBS, they were treated with fluorophore-labeled secondary antibodies, followed by washing with PBS.

Primary antibodies used were rabbit anti-actin polyclonal (Cytoskeleton, Denver, CO) and mouse anti–MYO-3 monoclonal (5–6; ref (65)) antibodies. Secondary antibodies used were Alexa 488–labeled goat anti-mouse IgG from Life Technologies and Cy3-labeled goat anti-mouse IgG from Jackson ImmunoResearch (West Grove, PA).

Fixed samples were mounted with ProLong Gold (Life Technologies) and observed by epifluorescence using a Nikon Eclipse TE2000 inverted microscope with a CFI Plan Fluor ELWD 40× (dry; NA 0.60) or Plan Apo 60× (oil; NA 1.40) objective. Images were captured by a SPOT RT monochrome charge-coupled device camera (Diagnostic Instruments, Sterling Heights, MI) and processed by IPLab imaging software (BD Biosciences) and Photoshop CS3 (Adobe, San Jose, CA) or by ORCA Flash 4.0 LT monochrome scientific complementary metal–oxide–semiconductor camera (Hamamatsu Photonics, Shizuoka, Japan) and processed by Nikon NIS-Elements and Adobe Photoshop CS3. Quantification of the myoepithelial sheath lengths was performed using ImageJ.

### Protein preparation

Actin was prepared from rabbit muscle acetone powder (Pel-Freeze Biologicals) as described by Pardee and Spudich (66). Recombinant LEV-11A (tropomyosin) (41), UNC-87B (22), and UNC-60B (ADF/cofilin) (45) were expressed in *Escherichia coli* and purified as described previously. Unstained molecular weight markers (Nacalai USA, catalog number 29458-24) were used in SDS-PAGE.

The open reading frame of *clik-1* cDNA was amplified with reverse-transcriptase-PCR and cloned at the NdeI - BamHI sites of pET-3a (Novagen) with no extra tag sequence. *E. coli* BL21(DE3) pLysS was transformed with the expression vector, cultured in M9ZB medium containing 50 μg/ml ampicillin and 34 μg/ml chloramphenicol at 37 °C until A600 reached 0.6 cm^−1^. Then, protein expression was induced by adding 0.1 mM isopropyl β-d-thiogalactopyranoside for 3 hr at 37 °C. The cells were harvested by centrifugation at 5000 × g for 10 min and disrupted by sonication in a buffer containing 0.1 M KCl, 1 mM EDTA, 20 mM Tris-HCl (pH 7.5), 0.2 mM dithiothreitol (DTT) and 1 mM phenylmethanesulfonyl fluoride. The homogenates were centrifuged at 25,000 × g for 30 min, and the supernatants were fractionated by 35 % saturated ammonium sulfate (194 g/l). Insoluble proteins were separated by centrifugation at 20,000 x g for 20 min, dissolved in a small volume of buffer A (0.1 M NaCl, 0.2 mM DTT, and 20 mM Tris-HCl, pH 7.5), and dialyzed overnight against buffer A. The dialyzed proteins were cleared by centrifugation at 20,000 x g for 20 min, applied to a Mono-Q 4.6/100PE column (GE Healthcare), and eluted with a linear gradient of NaCl from 0.1 to 0.5 M. Fractions containing CLIK-1 were applied to HiLoad 16/60 Superdex 75 (GE Healthcare) and eluted with a buffer containing 0.1 M KCl, 0.2 mM DTT, 20 mM HEPES-KOH, pH 7.5. Pure CLIK-1 fractions were pooled and concentrated by a MacroSep Centrifugal Device (molecular weight cutoff at 10K) (Pall Corporation), and stored at −80 °C. Protein concentrations were determined by a BCA Protein Assay kit (Thermo Scientific).

### F-actin sedimentation assays

Filamentous actin (5 or 10 μM) was incubated with proteins of interest in F-buffer (0.1 M KCl, 2 mM MgCl2, 20 mM HEPES-KOH, pH 7.5, 1 mM DTT) at room temperature. To examine F-actin binding of the proteins of interest by high-speed co-sedimentation, the samples were ultracentrifuged at 42,000 rpm (200,000 x g) for 20 min using a Beckman 42.2Ti rotor. To examine F-actin bundling by low-speed sedimentation, the samples were centrifuged at 15,000 rpm (18,000 x g) for 10 min using a Hettich MIKRO 200 centrifuge. Supernatants and pellets were separated, adjusted to the same volumes, and examined by SDS–PAGE (12% acrylamide gel). For quantitative analysis of high-speed F-actin cosedimentation, control experiments were performed without actin, and non-specific sedimentation of CLIK-1 was determined and subtracted from sedimentation of CLIK-1 in the presence of actin. The gels were stained with Coomassie Brilliant Blue R-250 (National Diagnostics) and scanned by an Epson Perfection V700 scanner at 300 dpi. Band intensity was quantified with ImageJ.

### Microscopic observation of actin bundles

Direct observation of actin bundles by fluorescent microscopy was performed as described previously (24,67) using DyLight 549-labeled actin (68). Briefly, DyLight 549-labeled actin filaments (2 μM actin, 20 % labeled) were incubated with proteins of interest for 5 min at room temperature in F-buffer, and 2 μl of the samples were applied to nitrocellulose-coated coverslips (22 x 40 mm) and mounted with noncoated coverslips (18 x 18 mm). They were immediately observed with epifluorescence using a Nikon Eclipse TE2000 inverted microscope with a Plan Apo 60× (oil; NA 1.40) objective. Images were captured by a SPOT RT monochrome charge-coupled device camera (Diagnostic Instruments, Sterling Heights, MI) and processed by IPLab imaging software (BD Biosciences) and Photoshop CS3 (Adobe, San Jose, CA). For quantification of actin bundles, regions of interest of 12.7 μm x 12.7 μm (100 x 100 pixels) were randomly selected. Then, using ImageJ, a threshold was set to eliminate single filaments, and number of pixels above the threshold regardless the intensity was counted.

### Statistical analysis

Data in Fig. 3I, 4Q, and 6K were analyzed by one-way analysis of variance with the Holm-Sidak method of pairwise multiple comparison procedures using SigmaPlot v13.0 (Systat Software, San Jose, CA).

### Data availability

All data are contained in the manuscript. Raw data are available from S. O. upon request.

## Acknowledgements

This paper is dedicated to the memory of Kanako Ono who passed away during the preparation of this manuscript.

## Author contribution

S.O. conceived the project and wrote the paper. S. O. and K. O. performed experiments, analyzed data, and prepared figures.

## References

1. Wu, K. C., and Jin, J. P. (2008) Calponin in non-muscle cells. Cell Biochem. Biophys. 52, 139–148

2. Carmichael, J. D., Winder, S. J., Walsh, M. P., and Kargacin, G. J. (1994) Calponin and smooth muscle regulation. Can. J. Physiol. Pharmacol. 72, 1415–1419

3. Winder, S. J., and Walsh, M. P. (1996) Calponin. Curr. Topics Cell. Reg. 34, 33–61

4. Gimona, M., Djinovic-Carugo, K., Kranewitter, W. J., and Winder, S. J. (2002) Functional plasticity of CH domains. FEBS Lett. 513, 98–106

5. Korenbaum, E., and Rivero, F. (2002) Calponin homology domains at a glance. J. Cell Sci. 115, 3543–3545

6. Gimona, M., and Mital, R. (1998) The single CH domain of calponin is neither sufficient nor necessary for F-actin binding. J. Cell Sci. 111, 1813–1821

7. Gimona, M., and Winder, S. J. (1998) Single calponin homology domains are not actin-binding domains. Curr. Biol. 8, R674–675

8. Galkin, V. E., Orlova, A., Fattoum, A., Walsh, M. P., and Egelman, E. H. (2006) The CH-domain of calponin does not determine the modes of calponin binding to F-actin. J. Mol. Biol. 359, 478–485

9. Stradal, T., Kranewitter, W., Winder, S. J., and Gimona, M. (1998) CH domains revisited. FEBS Lett. 431, 134–137

10. Gimona, M., Kaverina, I., Resch, G. P., Vignal, E., and Burgstaller, G. (2003) Calponin repeats regulate actin filament stability and formation of podosomes in smooth muscle cells. Mol. Biol. Cell 14, 2482–2491

11. Martin, R. M., Chilton, N. B., Lightowlers, M. W., and Gasser, R. B. (1997) *Echinococcus granulosus* myophilin--relationship with protein homologues containing “calponin-motifs”. Int. J. Parasit. 27, 1561–1567

12. Goodman, A., Goode, B. L., Matsudaira, P., and Fink, G. R. (2003) The Saccharomyces cerevisiae calponin/transgelin homolog Scp1 functions with fimbrin to regulate stability and organization of the actin cytoskeleton. Mol. Biol. Cell 14, 2617–2629

13. Winder, S. J., Jess, T., and Ayscough, K. R. (2003) SCP1 encodes an actin-bundling protein in yeast. Biochem. J. 375, 287–295

14. Nakano, K., Bunai, F., and Numata, O. (2005) Stg 1 is a novel SM22/transgelin-like actin-modulating protein in fission yeast. FEBS Lett. 579, 6311–6316

15. Pearlstone, J. R., Weber, M., Lees-Miller, J. P., Carpenter, M. R., and Smillie, L. B. (1987) Amino acid sequence of chicken gizzard smooth muscle SM22 alpha. J. Biol. Chem. 262, 5985–5991

16. Prinjha, R. K., Shapland, C. E., Hsuan, J. J., Totty, N. F., Mason, I. J., and Lawson, D. (1994) Cloning and sequencing of cDNAs encoding the actin cross-linking protein transgelin defines a new family of actin-associated proteins. Cell Motil. Cytoskeleton 28, 243–255

17. Takahashi, K., and Nadal-Ginard, B. (1991) Molecular cloning and sequence analysis of smooth muscle calponin. J. Biol. Chem. 266, 13284–13288

18. Yang, W., Zheng, Y. Z., Jones, M. K., and McManus, D. P. (1999) Molecular characterization of a calponin-like protein from *Schistosoma japonicum*. Mol. Biochem. Parasit. 98, 225–237

19. Matusovsky, O. S., Dobrzhanskaya, A. V., Pankova, V. V., Kiselev, K. V., Girich, U. V., and Shelud’ko, N. S. (2017) *Crenomytilus grayanus* 40kDa calponin-like protein: cDNA cloning, sequence analysis, tissue expression, and post-translational modifications. Comp. Biochem. Physiol. Part D, Genomics & Proteomics 22, 98–108

20. Wang, J., Gao, J., Xie, J., Zheng, X., Yan, Y., Li, S., Xie, L., and Zhang, R. (2016) Cloning and mineralization-related functions of the calponin gene in *Chlamys farreri. Comp*. Biochem. Physiol. Part B, Biochemistry & Molecular Biology 201, 53–58

21. Goetinck, S., and Waterston, R. H. (1994) The *Caenorhabditis elegans* muscle-affecting gene *unc-87* encodes a novel thin filament-associated protein. J. Cell Biol. 127, 79–93

22. Kranewitter, W. J., Ylanne, J., and Gimona, M. (2001) UNC-87 is an actin-bundling protein. J. Biol. Chem. 276, 6306–6312.

23. Lener, T., Burgstaller, G., and Gimona, M. (2004) The role of calponin in the gene profile of metastatic cells: inhibition of metastatic cell motility by multiple calponin repeats. FEBS Lett. 556, 221–226

24. Ono, K., Obinata, T., Yamashiro, S., Liu, Z., and Ono, S. (2015) UNC-87 isoforms, *Caenorhabditis elegans* calponin-related proteins, interact with both actin and myosin and regulate actomyosin contractility. Mol. Biol. Cell 26, 1687–1698

25. Yamashiro, S., Gimona, M., and Ono, S. (2007) UNC-87, a calponin-related protein in *C. elegans*, antagonizes ADF/cofilin-mediated actin filament dynamics. J. Cell Sci. 120, 3022–3033

26. Goetinck, S., and Waterston, R. H. (1994) The *Caenorhabditis elegans* UNC-87 protein is essential for maintenance, but not assembly, of bodywall muscle. J. Cell Biol. 127, 71–78

27. Waterston, R. H., Thomson, J. N., and Brenner, S. (1980) Mutants with altered muscle structure of *Caenorhabditis elegans*. Dev. Biol. 77, 271–302

28. Joseph, S., Gheysen, G., and Subramaniam, K. (2012) RNA interference in Pratylenchus coffeae: knock down of *Pc-pat-10* and *Pc-unc-87* impedes migration. Mol. Biochem. Parasit. 186, 51–59

29. Tan, J. A., Jones, M. G., and Fosu-Nyarko, J. (2013) Gene silencing in root lesion nematodes (*Pratylenchus spp.*) significantly reduces reproduction in a plant host. Exp. Parasit. 133, 166–178

30. Yuet, K. P., Doma, M. K., Ngo, J. T., Sweredoski, M. J., Graham, R. L., Moradian, A., Hess, S., Schuman, E. M., Sternberg, P. W., and Tirrell, D. A. (2015) Cell-specific proteomic analysis in *Caenorhabditis elegans*. Proc. Natl. Acad. Sci. U A 112, 2705–2710

31. Wang, H., Park, H., Liu, J., and Sternberg, P. W. (2018) An efficient genome editing strategy to generate putative null mutants in *Caenorhabditis elegans* using CRISPR/Cas9. G3 8, 3607–3616

32. McCarter, J., Bartlett, B., Dang, T., and Schedl, T. (1997) Soma-germ cell interactions in *Caenorhabditis elegans:* multiple events of hermaphrodite germline development require the somatic sheath and spermathecal lineages. Dev. Biol. 181, 121–143.

33. McCarter, J., Bartlett, B., Dang, T., and Schedl, T. (1999) On the control of oocyte meiotic maturation and ovulation in *Caenorhabditis elegans*. Dev. Biol. 205, 111–128.

34. Ardizzi, J. P., and Epstein, H. F. (1987) Immunochemical localization of myosin heavy chain isoforms and paramyosin in developmentally and structurally diverse muscle cell types of the nematode *Caenorhabditis elegans*. J. Cell Biol. 105, 2763–2770.

35. Strome, S. (1986) Fluorescence visualization of the distribution of microfilaments in gonads and early embryos of the nematode *Caenorhabditis elegans*. J. Cell Biol. 103, 2241–2252.

36. Ono, K., Yu, R., and Ono, S. (2007) Structural components of the nonstriated contractile apparatuses in the *Caenorhabditis elegans* gonadal myoepithelial sheath and their essential roles for ovulation. Dev. Dyn. 236, 1093–1105

37. Ono, K., and Ono, S. (2016) Two distinct myosin II populations coordinate ovulatory contraction of the myoepithelial sheath in the *Caenorhabditis elegans* somatic gonad. Mol. Biol. Cell 27, 1131–1142

38. Iwasaki, K., McCarter, J., Francis, R., and Schedl, T. (1996) *emo-1*, a *Caenorhabditis elegans* Sec61p gamma homologue, is required for oocyte development and ovulation. J. Cell Biol. 134, 699–714.

39. Kagawa, H., Sugimoto, K., Matsumoto, H., Inoue, T., Imadzu, H., Takuwa, K., and Sakube, Y. (1995) Genome structure, mapping and expression of the tropomyosin gene *tmy-1* of *Caenorhabditis elegans*. J. Mol. Biol. 251, 603–613.

40. Anyanful, A., Sakube, Y., Takuwa, K., and Kagawa, H. (2001) The third and fourth tropomyosin isoforms of *Caenorhabditis elegans* are expressed in the pharynx and intestines and are essential for development and morphology. J. Mol. Biol. 313, 525–537

41. Barnes, D. E., Watabe, E., Ono, K., Kwak, E., Kuroyanagi, H., and Ono, S. (2018) Tropomyosin isoforms differentially affect muscle contractility in the head and body regions of the nematode *Caenorhabditis elegans*. Mol. Biol. Cell 29, 1075–1088

42. Watabe, E., Ono, S., and Kuroyanagi, H. (2018) Alternative splicing of the *Caenorhabditis elegans lev-11* tropomyosin gene is regulated in a tissue-specific manner. Cytoskeleton (Hoboken) 75, 427–436

43. Ono, S., Baillie, D. L., and Benian, G. M. (1999) UNC-60B, an ADF/cofilin family protein, is required for proper assembly of actin into myofibrils in *Caenorhabditis elegans* body wall muscle. J. Cell Biol. 145, 491–502

44. McKim, K. S., Matheson, C., Marra, M. A., Wakarchuk, M. F., and Baillie, D. L. (1994) The *Caenorhabditis elegans unc-60* gene encodes proteins homologous to a family of actin-binding proteins. Mol Gen Genet 242, 346–357

45. Ono, S., and Benian, G. M. (1998) Two *Caenorhabditis elegans* actin depolymerizing factor/cofilin proteins, encoded by the *unc-60* gene, differentially regulate actin filament dynamics. J. Biol. Chem. 273, 3778–3783

46. Yamashiro, S., Mohri, K., and Ono, S. (2005) The two *Caenorhabditis elegans* actin depolymerizing factor/cofilin proteins differently enhance actin filament severing and depolymerization. Biochemistry 44, 14238–14247

47. Ono, K., and Ono, S. (2004) Tropomyosin and troponin are required for ovarian contraction in the *Caenorhabditis elegans* reproductive system. Mol. Biol. Cell 15, 2782–2793

48. Obinata, T., Ono, K., and Ono, S. (2010) Troponin I controls ovulatory contraction of nonstriated actomyosin networks in the *C. elegans* somatic gonad. J. Cell Sci. 123, 1557–1566

49. Myers, C. D., Goh, P. Y., Allen, T. S., Bucher, E. A., and Bogaert, T. (1996) Developmental genetic analysis of troponin T mutations in striated and nonstriated muscle cells of *Caenorhabditis elegans*. J. Cell Biol. 132, 1061–1077.

50. Ono, K., and Ono, S. (2014) Two actin-interacting protein 1 isoforms function redundantly in the somatic gonad and are essential for reproduction in *Caenorhabditis elegans*. Cytoskeleton 71, 36–45

51. Ono, K., Yamashiro, S., and Ono, S. (2008) Essential role of ADF/cofilin for assembly of contractile actin networks in the *C. elegans* somatic gonad. J. Cell Sci. 121, 2662–2670

52. Jensen, M. H., Morris, E. J., Gallant, C. M., Morgan, K. G., Weitz, D. A., and Moore, J. R. (2014) Mechanism of calponin stabilization of cross-linked actin networks. Biophys. J. 106, 793–800

53. Jensen, M. H., Watt, J., Hodgkinson, J. L., Gallant, C., Appel, S., El-Mezgueldi, M., Angelini, T. E., Morgan, K. G., Lehman, W., and Moore, J. R. (2012) Effects of basic calponin on the flexural mechanics and stability of F-actin. Cytoskeleton (Hoboken) 69, 49–58

54. Kolakowski, J., Makuch, R., Stepkowski, D., and Dabrowska, R. (1995) Interaction of calponin with actin and its functional implications. Biochem. J. 306, 199–204

55. Lu, F. W., Freedman, M. V., and Chalovich, J. M. (1995) Characterization of calponin binding to actin. Biochemistry 34, 11864–11871

56. Ciuba, K., Hawkes, W., Tojkander, S., Kogan, K., Engel, U., Iskratsch, T., and Lappalainen, P. (2018) Calponin-3 is critical for coordinated contractility of actin stress fibers. Sci. Rep. 8, 17670

57. Shapland, C., Hsuan, J. J., Totty, N. F., and Lawson, D. (1993) Purification and properties of transgelin: a transformation and shape change sensitive actin-gelling protein. J. Cell Biol. 121, 1065–1073

58. Han, M., Dong, L. H., Zheng, B., Shi, J. H., Wen, J. K., and Cheng, Y. (2009) Smooth muscle 22 alpha maintains the differentiated phenotype of vascular smooth muscle cells by inducing filamentous actin bundling. Life Sci. 84, 394–401

59. Gheorghe, D. M., Aghamohammadzadeh, S., Smaczynska-de, R., II, Allwood, E. G., Winder, S. J., and Ayscough, K. R. (2008) Interactions between the yeast SM22 homologue Scp1 and actin demonstrate the importance of actin bundling in endocytosis. J. Biol. Chem. 283, 15037–15046

60. Stiernagle, T. (2006) Maintenance of *C. elegans*. WormBook, 1–11

61. Dickinson, D. J., Pani, A. M., Heppert, J. K., Higgins, C. D., and Goldstein, B. (2015) Streamlined genome engineering with a self-excising drug selection cassette. Genetics 200, 1035–1049

62. Ono, S., and Ono, K. (2002) Tropomyosin inhibits ADF/cofilin-dependent actin filament dynamics. J. Cell Biol. 156, 1065–1076.

63. Timmons, L., Court, D. L., and Fire, A. (2001) Ingestion of bacterially expressed dsRNAs can produce specific and potent genetic interference in *Caenorhabditis elegans*. Gene 263, 103–112.

64. Ono, S. (2001) The *Caenorhabditis elegans unc-78* gene encodes a homologue of actin-interacting protein 1 required for organized assembly of muscle actin filaments. J. Cell Biol. 152, 1313–1319.

65. Miller, D. M., Ortiz, I., Berliner, G. C., and Epstein, H. F. (1983) Differential localization of two myosins within nematode thick filaments. Cell 34, 477–490

66. Pardee, J. D., and Spudich, J. A. (1982) Purification of muscle actin. Methods Enzymol. 85, 164–181

67. Ono, S. (2016) Basic Methods to Visualize Actin Filaments In Vitro Using Fluorescence Microscopy for Observation of Filament Severing and Bundling. in Cytoskeleton: Methods and Protocols, 3rd Edition (Gavin, R. H. ed.). pp 187–193

68. Liu, Z., Klaavuniemi, T., and Ono, S. (2010) Distinct roles of four gelsolin-like domains of *Caenorhabditis elegans* gelsolin-like protein-1 in actin filament severing, barbed end capping, and phosphoinositide binding. Biochemistry 49, 4349–4360

